# Unraveling the Dual-Mode Impact on Tension Gauge Tethers’ Mechanical Stability

**DOI:** 10.1101/2023.08.10.552888

**Authors:** Jingzhun Liu, Jie Yan

## Abstract

Tension Gauge Tethers (TGTs), short DNA segments serving as extracellular tension sensors, are instrumental in assessing tension dynamics in mechanotransduction. These TGTs feature an initial shear-stretch region and an unzip-stretch region. Despite their utility, no theoretical model has been available to estimate their tension-dependent lifetimes (*τ* (*f*)), restricting insights from cellular mechanotransduction experiments. We’ve now formulated a concise expression for *τ* (*f*) of TGTs, accommodating contributions from both stretch regions. Our model uncovers a tension-dependent energy barrier shift occurring when tension surpasses a switching force approximately 13 pN for the recently developed TGTs, greatly influencing *τ* (*f*) profiles. Experimental data from several TGTs validated our model. The calibrated expression can predict *τ* (*f*) of TGTs at 37 degrees Celsius based on their sequences with minor fold-changes, supporting future applications of TGTs.

---

Tension Gauge Tethers (TGTs) are short segments of double-stranded DNA (dsDNA) that irreversibly dissociate in response to tension. TGTs were initially developed as sensors to measure tension levels transmitted across tension-bearing extracellular receptors, such as integrin, cadherin, and T-cell receptors [1–8]. In these applications, a TGT is incorporated into a tension-transmission pathway via two attachment points on the TGT, which consists of a total of *N* base pairs. The first point, *P*_1_, is located at the end of one single-stranded DNA (ssDNA) strand, and the second point, *P*_2_, is situated on the opposite strand, *M* base pairs away from *P*_1_ (see (Fig. 1a)). Therefore, the stretching geometry of a TGT with a specific sequence is entirely defined by *N* and *M*, and is henceforth denoted as M/N-TGT ((Fig. 1b)).

**Fig. 1.**
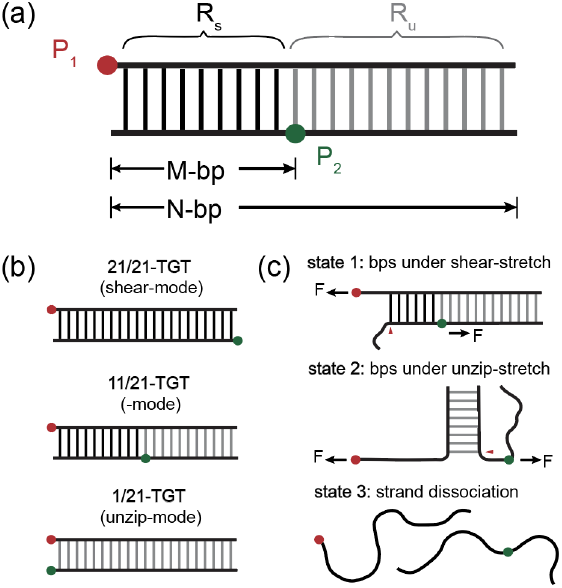
Illustration of TGTs and the transition pathway for tension-dependent strand dissociation pathway. (a) *P*_1_ (red) and *P*_2_ (green) are two force attaching point on a TGT with a total length of *N* -bp that divides it into two regions: *R*_s_ (black) and *R*_u_ (grey) under shear- and unzip-stretching geometry, respectively. A TGT is named as M/N-TGT where *M* is the base pair distance between *P*_1_ and *P*_2_. (b) Typical design of TGTs with *P*_2_ at different positions (from top to bottom: left-end, middle, and right-end) on DNA strand, named as 21/21-bp, 11/21-bp, and 1/21-bp TGT. (c) The tension-dependent strand dissociation pathway of a TGT via strand peeling from *P*_1_ end.

A TGT can be divided into two distinct regions: *R*_s_, which exists between *P*_1_ and *P*_2_ under a shear-stretching geometry, and *R*_u_, which exists after *P*_2_ under an unzip-stretching geometry (as shown in Fig. 1c). The strand dissociation of a TGT, which depends on the tension, follows a lowest-energy kinetic pathway. This pathway involves sequential breakage of base pairs from the *P*_1_ end until the strands are fully dissociated (as shown in Fig. 1c). The transition state can be indicated by the number of base pairs (*n*) between the fork and *P*_1_. It is clear that there is a switch in stretching geometry when the fork passes *n* = *M −* 1.

Our current understanding of the mechanical stability of TGTs remains incomplete. The original description of mechanical stability have been based on the tension tolerance, *T*_tol_, which is the tension required to dissociate a TGT with a given average lifetime of 2 seconds as per the studies by Wang et al. [1]. Recent research underscores the significance of comprehending the tension-dependent lifetime, *τ* (*f*) [9], which offers a comprehensive understanding of the mechanical stability of TGTs. Despite its importance, no theoretical model has been available to estimate *τ* (*f*) for TGTs based on their sequences and stretching geometry, restricting insights from cellular mechanotransduction experiments.

In this study, we utilized Kramers’ kinetic theory to derive a general expression for the TGTs’ *τ* (*f*), which has a single model parameter, the effective diffusion coefficient *D* of DNA fork migration (depicted by red arrows in Fig. 1c) during the transition. Our expression depends on the tension-dependent, one-dimensional energy landscape, which is influenced by various factors, including the DNA sequence, the entropic elasticity of both double-stranded (ds) and single-stranded (ss) DNA, and the stretching geometries involved.

The process of strand dissociation of a TGT can be understood by considering the random movement of the fork position, with the rates of base pair opening (forward) and formation (backward) depending on the tension. The energy required to open the *i*^*th*^ base pair from the ends can be expressed as:

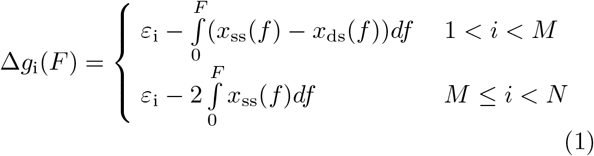

Here *x*_ss_(*f*) and *x*_ds_(*f*) are the force-extension curves for 1-nt ssDNA and 1-bp dsDNA, respectively, and *ε*_i_ represents the energy cost to open the base pair at zero tension, which is evaluated in the work using the Santa Lucia nearest-neighbour base pair energy data [10].

We calculated tension-dependent energy landscapes for several commonly used TGTs with a given sequence, *N*, and *M* (see Fig. 2). For forces below a threshold of 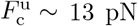, the energy profile monotonically increases with the number of opened base pairs until the second last base pair in the TGT is ruptured. However, at higher forces, the energy profile is non-monotonic, with a maximum at the base pair position *n* = *M−*1. Our analysis suggests a tension-dependent switch in the energy barrier. Note that the threshold tension 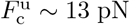, is critical because it represents the tension at which the opening and closing rates of a base pair are equal under the unzip-stretching geometry.

**Fig. 2.**
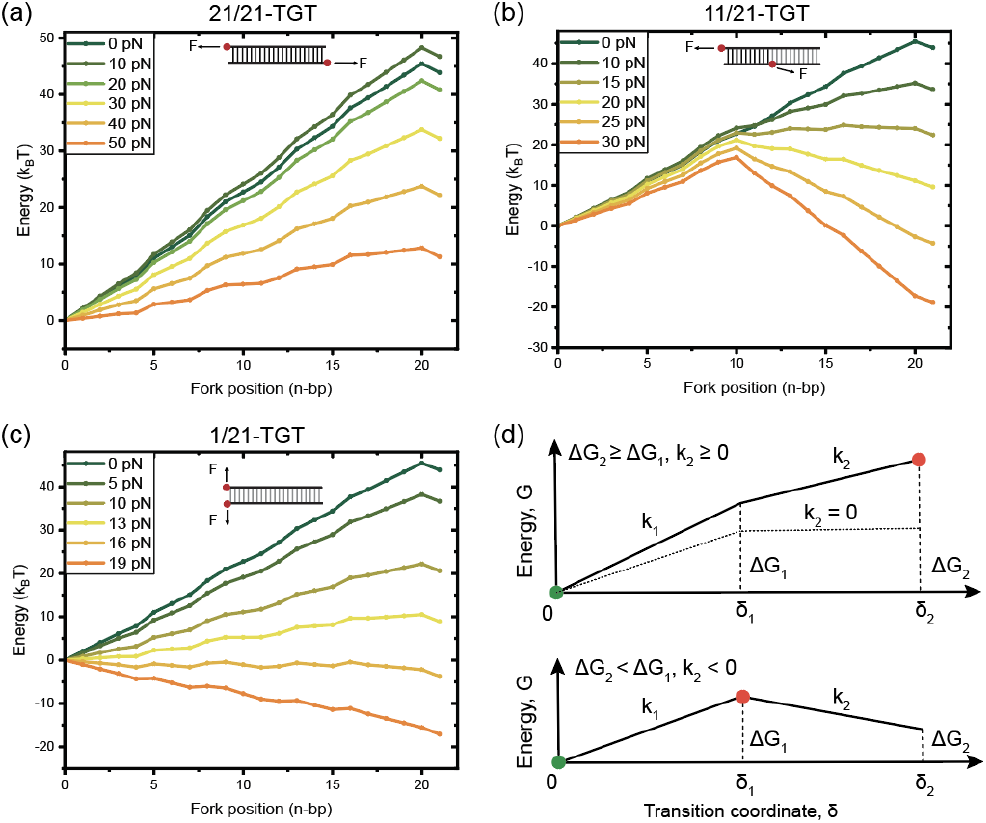
Tension-dependent energy landscape of a series of 21-bp TGT with different stretching modes (5’-GTGTCGTGCCTCCGTGCTGTG-3’): (a) 21/21-TGT in shear-stretch mode, (b) 11/21-TGT in dual-mode, and (c) 1/21-TGT in unzip-stretch mode. The transition distance, the x-axis, is indicated by the fork position *n*. The tension-dependent energy landscapes of these TGTs are plotted over a wide force range from zero to a force that the dissociation state have much lower free energy than the hybridized state. (d) Illustration of the simplified energy landscapes of the 11/21-TGT with 10-bp in shear-mode and 11-bp in unzip-mode.

It can be seen from Figure 2 that for TGTs with the sequence 5’-GTGTCGTGCCTCCGTGCTGTG-3’ (*N* = 21), at a given tension, the energy landscape can be approximated by a piecewise linear function:

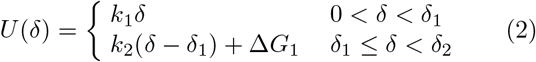

Here, *δ* is a force-independent transition coordinate, chosen to be the number of dissociated base pairs during transition. The energy landscape is composed of two straight lines with different slopes, each defined by four geometric parameters. The first set of parameters, *δ*_1_ and Δ*G*_1_, describe the number of dissociated base pairs and energy difference between the fully hybridized state (*n* = 1) and the state at fork position *n* = *M −* 1. The second set of parameters, *δ*_2_ and Δ*G*_2_, describe the difference between the fully hybridized state (*n* = 1) and the rupture of the second last base pair(*n* = *N−* 1) which breaks the stacking energy of the last base pair step (as shown in Fig. 2d).

The four parameters are not independent variables; rather, they are directly derived from the values of *M, N*, and the tension-dependent base pair energy data Δ*g*_i_. Specifically, they are calculated as follows: 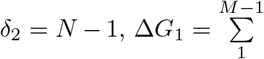 and 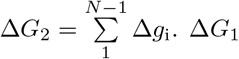 and Δ*G*_2_ are normalized into a dimensionless quantity by dividing by *k*_B_*T*, where *k*_B_ is the Boltzmann constant and *T* is the temperature. The slopes of the two lines are determined by the corresponding parameters: 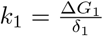 and 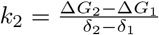 respectively.

We examine *τ* (*f*) of a TGT, *k*(*f*), based on the energy landscape Eq. 2 within the framework Kramer’s kinetic theory [11]. Following the derivation reported previously [12, 13], it can be shown that (SI-III):

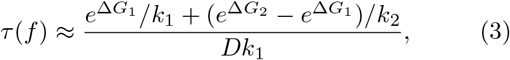

where the single model parameter *D* is the effective diffusion coefficient for fork migration during the transition process. The above formula has assumed sufficiently high value Δ*G*_1_ to meet the pre-equilibration condition imposed by the Kramer’s kinetic theoretical framework [11].

At tensions much greater than 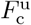, Δ*G*_1_ ≫ Δ*G*_2_ and so 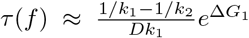, indicating that the lifetime is determined by the barrier Δ*G*_1_ . On the contrary, at tensions much smaller than 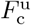, Δ*G*_1_ ≪ Δ*G*_2_ and so 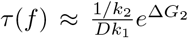, indicating that the lifetime is determined by Δ*G*_2 ._ The result indicates a switch of the energy barrier from Δ*G*_1_ to Δ*G*_2_ when the tension decreases from below to above 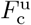.

To evaluate the predictive capabilities of the model for dual-mode TGTs containing both shear-stretch and unzip-stretch regions at various forces, we conducted experiments for five TGTs, 11/11-TGT, 11/16-TGT, 11/21-TGT, 15/21-TGT, and 20/21-TGT (Fig. 3a). The first three TGTs share the same shear-stretch region, but differ in the length of their unzip-stretch regions, while the last three TGTs have the same overall length but feature different separations of shear-stretch and unzip-stretch regions (Fig. 3b).

**Fig. 3.**
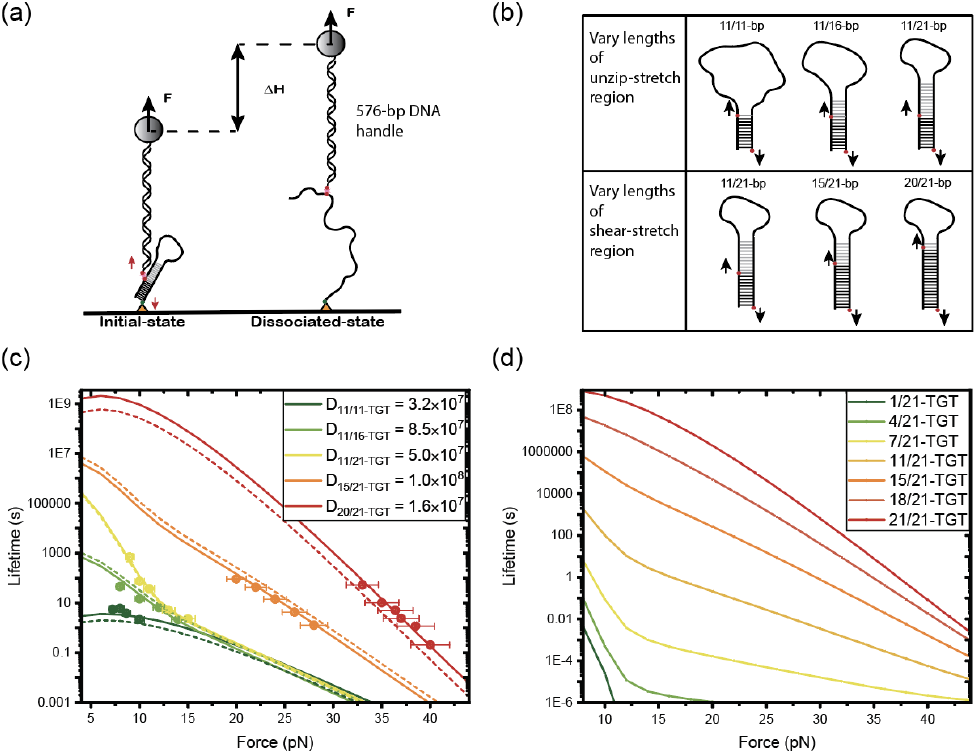
Single-molecule quantification of *τ* (*f*) of five different TGTs and the prediction of *τ* (*f*) of currently widely utilized TGTs (N = 21). (a) Schematic of the designed DNA detector tethered between a glass surface and a superparamagnetic bead. The TGT is stretched through a thiol anchored on glass surface and a biotin linked to a 576-bp DNA handle attached on the superparamagnetic bead. (b) The stretching geometry of our examined TGTs with varying lengths in unzip- or shear-stretch regions. (c) *τ* (*f*) of 11/11-, 11/16-, 11/21-TGT, 15/21-, and 20/21-TGT. The solid lines represent the best-fitting curves for the quantified data points, while the dashed lines depict the *τ* (*f*) values predicted by the average of the five best-fitting parameters *D*, obtained from the five TGTs. (d) The predicted *τ* (*f*) of seven widely utilized TGTs in cellular experiment (N = 21).

For each TGT, the average lifetimes were determined by analyzing at least 30 force-jump cycles for each target tension, using data from three individual detectors. One exception is the data point obtained at 9 pN for the 11/21 TGT, where only nine data points were obtained from a single tether, due to the long lifetime at this force. As tension increases, the lifetime of all these TGTs decreases, indicating that they become less stable under higher tension. As depicted in Fig. 3c (solid lines), the model can reasonably fit the data obtained from all the TGTs, with fitting values of *D* over a range from 1.6 *×* 10^7^ s^*−*1^ to 1.0 *×* 10^8^ s^*−*1^, by introducing a small constant energy correction *σ* = 0.45 *k*_B_*T* to account for any inaccuracy in tension-dependent base pair energy (e.g., the force-extension of ssDNA at the diffusing fork (SI-IV). Using an average value of approximately *D* = 5.7*×* 10^7^ s^*−*1^, the predicted *τ* (*f*) agrees reasonably well with all the indiviudally fitted profiles of these experimentally quantified TGTs (Fig. 3c, dashlines with conresponding colors).

As the contour length of a dsDNA base pair is *l*_ds_ = 0.34 nm, the effective diffusion coefficient *D* can be converted to 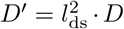 with the unit of nm^2^ *·* s^-1^.The value of *D*^*I*^ is over the range from 1.9 *×* 10^6^ nm^2^ s^*−*1^ to 1.2*×* 10^7^ nm^2^ s^*−*1^, which is similar to the magnitude reported in previous studies [14, 15].

Collectively, these findings indicate that the model (Eq. 3) can be applied to various TGTs with differing lengths, diverse stretch region-to-unzip region ratios, and both lower 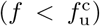 and higher 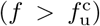 forces. Although the fork diffusion constant *D* fitting values vary among TGTs, the fold differences remain within one order of magnitude. These disparities in the fitting values of *D* can be attributed to inaccuracies in estimating tension-dependent base pair energies (SI-IV).

By utilizing the average value of *D*, we were able to estimate *τ* (*f*) for seven 21-bp TGTs that were investigated in previous cell experiments [1–8] (as depicted in Figure 3d). The plot reveals highly different tension-dependent lifetime profiles due to the different positions of *P*_2_. When *P*_2_ is located near the middle of the DNA segment, the profile shows a clear change of slope slightly above the value of 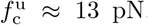. Since the highly diverse *τ* (*f*) profiles result from changes in *P*_2_ that alter the lengths of the shear-stretch and unzip-stretch regions, this strongly suggests differential contributions from both regions to the tension-dependent lifetime of TGTs.

Overall, the findings suggest that, at high forces 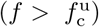, the transition rate is primarily influenced by the energy barrier’s height at position *M −* 1. Additionally, the unzip-stretch region has a marginal impact on the transition rate at forces slightly higher than 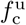. At lower forces 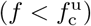, both the shear and unzip-stretch regions contribute to the tension-dependent lifetime *τ* (*f*). The significantly increased lifetime in the lower force range is a direct result of the higher energy cost of breaking each base pair, the switched energy barrier, and the longer diffusion distance. Further details on the differential contributions from shear- and unzip-stretch regions at low or high force regions are provided in Supplementary Informations (SI-V and VI).

Tension Gauge Tethers (TGTs) offer comprehensive insights into the mechanical responses of mechanosensing systems and tension-transmission in supramolecular linkages. Beyond their originally proposed role as tension-threshold sensors, requiring a lifetime of a few seconds, TGTs can serve a dual function during dynamic stretching processes. Specifically, TGTs can indicate whether the applied tension surpasses the threshold needed to activate key mechanosensing proteins, and whether the duration of this tension is sufficient for the activated proteins to facilitate downstream mechanotransduction processes.

To substantiate this, Figure 4 displays the rupture forces (Figure 4a) and lifetimes (Figure 4b) of seven commonly used TGTs across various loading rates. Consider the 11/21-TGT and 15/21-TGT as examples: While both have been demonstrated to support cell adhesion spreading, cells spread on the 15/21-TGT surface exhibit greater spreading and focal adhesion areas than those on the 11/21-TGT. Figure 4a reveals that at loading rates of *>* 1 pN/s, both TGTs rupture at forces *>* 10 pN, a magnitude exceeding the activation forces required for talin mechanosensing domains via mechanical unfolding [9**?**]. Furthermore, Figure 4b demonstrates that the lifetime of the 15/21-TGT is significantly longer than that of the 11/21-TGT at identical loading rates. This potentially allows for an extended duration of downstream mechanotransduction processes, offering a plausible mechanism to explain the distinct cell adhesion behaviors observed on surfaces coated with these specific TGTs.

**Fig. 4.**
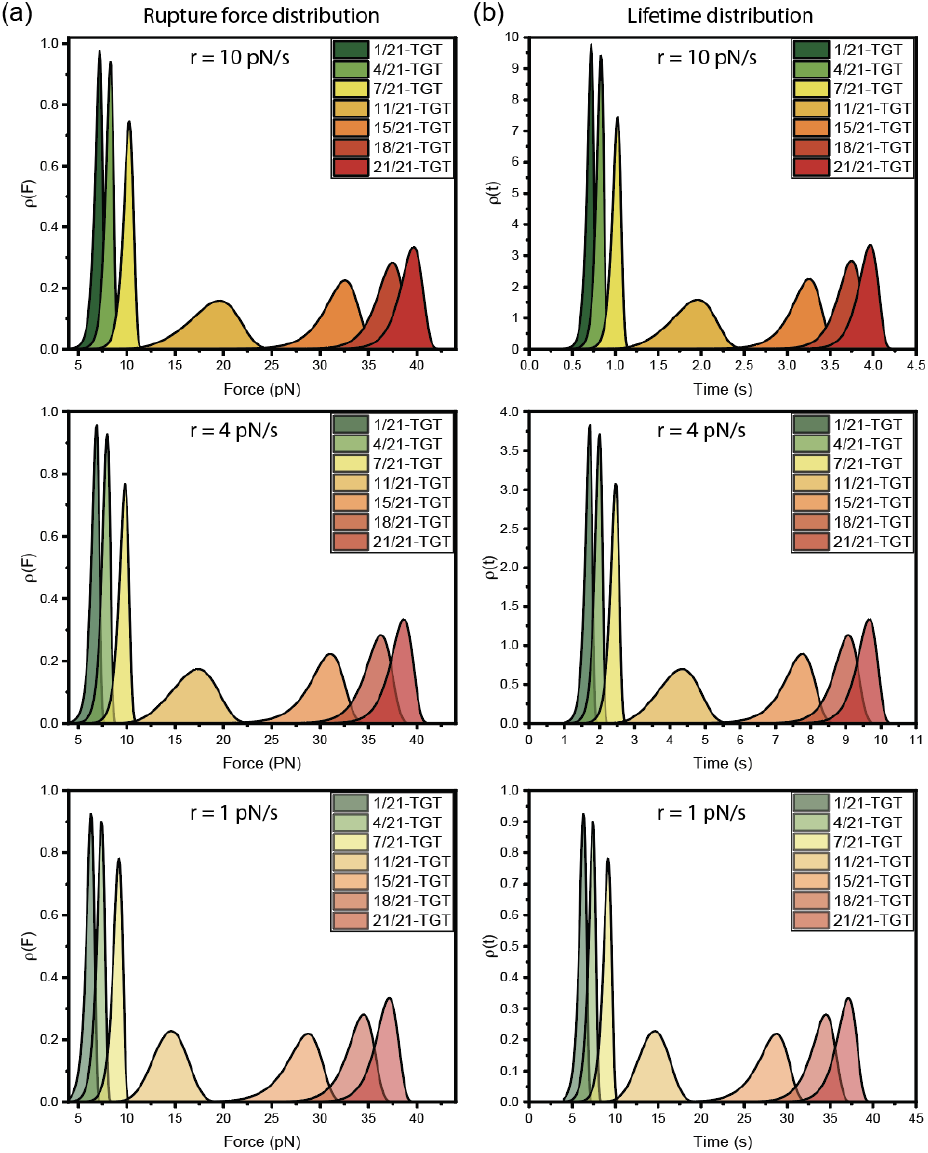
The rupture force distributions (a) and the life-time distributions (b) of seven widely utilized TGTs (N = 21) with different force increasing rates derived from the predicted *τ* (*f*) of these TGTs shown in Fig. 3d. From top to bottom panel, the increasing rate are r = 10 pN/s, 4 pN/s, and 1 pN/s, respectively.

In summary, through the analysis of the energy landscape of dual-mode TGTs and the application of Kramer’s kinetic theory, we have derived a simple analytical expression, *τ* (*f*), for which the fork diffusion constant serves as the single model parameter. The energy parameters, Δ*G*_1_ and Δ*G*_2_, are calculated from the DNA sequences using the Santa Lucia nearest-neighbour base pair energy data and the force-extension curves of the ds and ssDNA. The expression was calibrated at a temperature of 37 °C, which aligns with the conditions of most cell experiments. The calibrated expression yielded a range of fork diffusion constants between 1.6 *×* 10^7^ s^*−*1^ and 1.0 *×* 10^8^ s^*−*1^ for five different TGTs with varying lengths and shear-to-unzip base pair ratios. Using the average of the diffusion coefficient, the expression can predict *τ* (*f*) with reasonable accuracy solely based on the sequence and the stretching geometry of TGTs, facilitating their use in cell mechanotransduction studies conducted at 37 degrees Celsius.

## ACKNOWLEDGEMENTS

The research was funded by the Singapore Ministry of Education Academic Research Funds Tier 2 (MOE-T2EP50220-0015), the Singapore Ministry of Education Academic Research Fund Tier 3 (MOE Grant No: MOET32021-0003) and the Ministry of Education under the Research Centres of Excellence programme.

