## Supplementary Information for "Unraveling the Dual-Mode Impact on Tension Gauge Tethers’ Mechanical Stability" (from another version of this preprint: version 1, 2023-08-14)

### Supplemental Material

#### I. Single-molecule constructs

All DNA oligos used in this study were custom synthesized by Integrated DNA Technologies (Coralville, Iowa) shown in Table SI. The DNA-based detector with two dsDNA handles and a hairpin region was synthesized following the standard PCR, digestion, annealing, ligation, and purification protocols.

#### II. Single-molecule manipulation

The single-molecule manipulation experiments were conducted using an in-house-made back-scattered vertical magnetic tweezers with a spatial resolution of  1 nm and temporal resolution of  200 Hz. DNA-based detector was tethered to the coverslip through Thiol/SMCC reaction, while the other end of the detector was linked to a superparamagnetic bead through Biotin/Traptavidin linkage. This system was performed in a laminar flow channel. The extension change of the tethered DNA was measured based on the height change of the superparamagnetic beads tethered to the DNA-based detector under force. The details of the force calibration and control for the single-molecule magnetic tweezers experiments have been described in previous papers [1, 2].

Experiments were performed in a standard solution containing: 1 × PBS and 1% BSA at 37 °C. To achieve temperatures higher than room temperature, a non-magnetic nanocarbon-based thermal film (Ultra-Thin Flexible Heater, Pelonis Technologies) was applied on the top of the sample channel.

#### III. Deviation for analytical expression of tension-dependent lifetime of TGTs

In this study, we derived a general expression for the tension dependent lifetime of strand-dissociation of a TGT, $\tau(f)$ based on the energy landscape within the framework Kramer's kinetic theory [3].

The diffusive flux under influence of potential from energy landscape has two contributions:

$$J(\xi) = -D\frac{dC(\delta)}{d\delta} - C(\delta)\zeta\frac{dU(\delta)}{d\delta} \tag{1}$$

where $C(\xi)$ is the concentration, $D$ is the diffusion coefficient, $\zeta$ is the friction coefficient, and $U(\xi)$ is the potential energy as a function of position $\delta$. The first term on the right-hand side of the equation represents the contribution

TABLE SI: DNA sequences. Underscore indicates the DNA segments that form the TGT region under shearing force geometry.

| Oligo ID | Sequence (5' → 3') |
|---|---|
| 11+0-bp_lifetime | GTGTCGTGCC/iBiodT/ATCTTTTATCCTATTCCTATCATTATTC ATTATTTCTACCAACCAGGCACGACACTCATCGACGTTTTTAT CCGGAAACTGCTGTCTGG/3ThioMC3-D/ |
| 11+5-bp_lifetime | GTGTCGTGCC/iBiodT/CCGTGTTATCCTATTCCTATCATTATT CATTATTTCTACCACGGAGGCACGACACTCATCGACGTTTTTA TCCGGAAACTGCTGTCTGG/3ThioMC3-D/ |
| 11+10-bp_lifetime | GTGTCGTGCC/iBiodT/CCGTGCTGTGCTATTCCTATCATTAT TCATTATTCACAGCACGGAGGCACGACACTCATCGACGTTTT TATCCGGAAACTGCTGTCTGG/3ThioMC3-D/ |
| 15+6-bp_lifetime | GTGTCGTGCCTCCG/iBiodT/GCTGTGTTTTTTTCACAG CACGGAGGCACGACACTTCATCGACGTTTTTATCCGGAAACT GCTGTCTGGCTTTTTTTGATTTCA/3ThioMC3-D/ |
| 20+1-bp_lifetime | GTGTCGTGCCTCCGTGCTG/iBiodT/GTATAATATTTTTTTTAT ATTATACACAGCACGGAGGCACGACACTTCATCGACGTTTTT ATCCGGAAACTGCTGTCTGG/3ThioMC3-D/ |
| Com_handle | CCAGACAGCAGTTTCCGGATAAAAACGTCGATG |

from the concentration gradient $\frac{dC(\delta)}{d\delta}$, which is proportional to the diffusion coefficient $D$. The second term represents the contribution from the force due to the gradient of potential $dU(\delta)/d\delta$.

As discussed earlier, $\zeta$ is the friction coefficient and in one dimension:

$$\zeta = \frac{k_B T}{D} \tag{2}$$

This can be written in terms of a probability density $P$:

$$J = D\left(-\frac{P}{k_B T}\frac{dU}{d\delta} - \frac{dP}{d\delta}\right) = -De^{-U(\delta)/k_B T}\frac{d}{d\delta}\left(Pe^{U(\delta)/k_B T}\right) \tag{3}$$

or

$$Je^{U(\delta)/k_B T} = -D\frac{d}{d\delta}\left(Pe^{U(\delta)/k_B T}\right) \tag{4}$$

Here, it is assumed that $D$ and $\zeta$ are not functions of $\delta$.

The transition rate $k$ can be written as:

$$k = -De^{-\beta U(\delta)}\frac{d}{d\delta}\left(\rho(\delta)e^{\beta U(\delta)}\right) \tag{5}$$

where $\beta = 1/k_B T$, and $\rho(\delta)$ is the probability density.

The next important assumption of Kramers' theory is that we can solve for the transition rate using the steady-state approximation. This allows us to set $J =$ constant. Integrate along $\delta$ over the energy landscape.

$$-\frac{k}{D}\int_{x_1}^{x_2} e^{\beta U(\delta)}d\delta = \int_{x_1}^{x_2}\frac{d}{d\delta}\left(\rho(\delta)e^{\beta U(\delta)}\right) \tag{6}$$

In the case of TGT strand dissociation, the integration upper limit $x_2 = \delta_2$ and the lower limit $x_1 \ll \delta_1$.

$$-\frac{k}{D}\left(\int_{x_1}^{\delta_1} e^{\beta U(\delta)}d\delta + \int_{\delta_1}^{\delta_2} e^{\beta U(\delta)}d\delta\right) = \rho(\delta_2)e^{\beta U(\delta_2)} - \rho(x_1)e^{\beta U(x_1)} \tag{7}$$

If diffusion across the $\delta_2$ boundary, the probability density will become zero, which means $\rho(\delta_2) = 0$. Then, assume a Boltzmann distribution at the energy well (from $-x_1$ to $x_1$, $x_1 \gg 0$), $\rho(x_1) = \frac{\beta e^{U(x_1)}}{Z_w}$, where $Z_w$ is the partition function that can be expressed as $Z_w = \int_0^{x_1} e^{-\beta U(\delta)}d\delta$. Then, the integration can be written as

$$-\frac{k}{D}\left(\int_{x_1}^{\delta_1} e^{\beta U(\delta)}d\delta + \int_{\delta_1}^{\delta_2} e^{\beta U(\delta)}d\delta\right) = -\frac{1}{Z_w} \tag{8}$$

$$k = \frac{D}{\int_0^{x_1} e^{-\beta U(\delta)}d\delta\left(\int_{x_1}^{\delta_1} e^{\beta U(\delta)}d\delta + \int_{\delta_1}^{\delta_2} e^{\beta U(\delta)}d\delta\right)} \tag{9}$$

This way, for the piece-wise linear shaped energy landscape described in this study, the tension-dependent transition rate can be simplified as following:

$$\begin{aligned} k(f) &= \frac{D}{\int_0^{x_1} e^{-\beta k_1 \delta}d\delta\left(\int_{x_1}^{\delta_1} e^{\beta k_1 \delta}d\delta + \int_{\delta_1}^{\delta_2} e^{\beta(k_2(\delta-\delta_1)+\Delta G_1)}d\delta\right)} \\ &= \frac{-D\beta k_1}{(e^{-\beta k_1 x_1} - 1)\left((e^{\beta k_1 \delta_1} - e^{\beta k_1 x_1})/\beta k_1 + e^{\beta \Delta G_1 - \beta k_2 \delta_1}(e^{\beta k_2 \delta_2} - e^{\beta k_2 \delta_1})/\beta k_2\right)} \\ &\approx \frac{D\beta^2 k_1}{e^{\beta \Delta G_1}/k_1 + (e^{\beta \Delta G_2} - e^{\beta \Delta G_1})/k_2} \end{aligned} \tag{10}$$

##### IV. The constant energy correction $\sigma$ introduced for model-fitting

As illustrated in the manuscript, the model adeptly accommodates the data derived from all five TGTs, with fitting values of $D$ spanning a range from $2.38 \times 10^{8}\ s^{-1}$ to $1.56 \times 10^{9}\ s^{-1}$. This achievement is facilitated by the incorporation of a minor constant energy correction, denoted as $\sigma = 0.25\ k_{\mathrm{B}}T$, aimed at compensating for potential inaccuracies in tension-dependent base pair energy. The introduction of this constant energy correction serves to rectify any imprecision that may arise during the computation of force-dependent base pair energies from the force-extension curve and during the execution of the piecewise linear approximation. For example, the worm-like chain model of ssDNA fails at forces below 8 pN [4], where ssDNA forms a more compact conformation due to random base stacking and hydrogenbonding [5]. Further, the value of $\sigma$ is expected to be dependent on the salt concentration as it affects both the zero-force base pair stability and the force-extension curve of ssDNA.

##### V. Differential contributions from shear- and unzip-stretch regions at $f > f_{\mathrm{c}}^{\mathrm{u}}$

As aforementioned, at forces greater than $f_{\mathrm{c}}^{\mathrm{u}}$, the energy barrier is located at the second last base pair in the shear-stretch region, suggesting a predominant contribution of this region to $\tau(f)$. However, it remains unclear regarding the potential contribution from the unzip-stretch region that provides a space for over-barrier diffusion of the fork. To understand the differential contributions from shear- and unzip-stretch regions at forces $> f_{\mathrm{c}}^{\mathrm{u}}$, we predicted the normalized tension-dependent lifetimes for several TGTs with controlled shear- and unzip-stretch regions.

Figure S1a shows the energy landscapes of three TGTs, 11/17-TGT, 13/19-TGT, 15/21-TGT, at a force of 20 pN. These TGTs have the same length of the unzip-stretch region but different lengths of the shear force region. It shows that with an increase in the length of the shear-stretch region by two base pairs, the energy barrier of the TGT increased by several $k_B T$. This is expected to increase the TGT's lifetime by 1 to 2 orders of magnitude, consistent with the predicted normalized tension-dependent lifetime at forces above $f_{\mathrm{c}}^{\mathrm{u}}$ (Fig. S1b).

Figure S1c presents the energy landscapes of three TGTs - 11/11-TGT, 11/16-TGT, and 11/21-TGT - at a force of 20 pN. Although these TGTs have the same shear-stretch region length, they possess different unzip-stretch region lengths. The figure demonstrates that just a few pN above $f_{\mathrm{c}}^{\mathrm{u}}$, the length of the unzip-stretch region significantly influences $\tau(f)$; a longer unzip-stretch region corresponds to a longer lifetime. However, at higher forces (above 20 pN), the impact of the unzip-stretch region diminishes, as evidenced by the nearly overlapping $\tau(f)$ profiles of the 11/16-TGT and 11/21-TGT (Fig. S1d).

Overall, the findings in this section suggest that at greater forces ($f > f_{\mathrm{c}}^{\mathrm{u}}$), the transition rate is predominantly constrained by the energy barrier's height at position $M - 1$. Additionally, the unzip-stretch region influences the transition rate at forces marginally higher than $f_{\mathrm{c}}^{\mathrm{u}}$.

##### VI. Differential contributions from shear- and unzip-stretch regions at $f < f_{\mathrm{c}}^{\mathrm{u}}$

We utilized the model to predict $\tau(f)$ for smaller forces $f < f_{\mathrm{c}}^{\mathrm{u}}$, where the energy barrier is shifted to $N - 1$ base pairs in the TGT. This results in a significantly higher energy barrier and a longer distance for the fork to diffuse, both of which contribute to an increased lifetime of the TGTs. To determine the differential contributions from the shear- and unzip-stretch regions, we predicted the normalized tension-dependent lifetimes for the TGTs with controlled shear- and unzip-stretch regions.

In Figure S2a, we present the energy landscapes of three TGTs, 11/17-TGT, 13/19-TGT, and 15/21-TGT, at a force of 10 pN. The length of the shear-stretch region was increased by two base pairs in each TGT, resulting in a several $k_B T$ increase in the energy barrier and a 1–2 order of magnitude increase in lifetime (Fig. S2b). The model also predicts significantly longer force-dependent lifetimes compared to those at higher forces $f > f_{\mathrm{c}}^{\mathrm{u}}$.

Figure S2c displays the energy landscapes of three TGTs, 11/11-TGT, 11/16-TGT, and 11/21-TGT, at a force of 10 pN. As the length of the unzip-stretch region increases, the energy barrier height increases, resulting in a major dependence of $\tau(f)$ on the length of the unzip-stretch region. When the length of the unzip-stretch region increases by 5 base pairs, the lifetime can increase by several orders of magnitude.

In summary of this section, our results indicate that at lower forces ($f < f_{\mathrm{c}}^{\mathrm{u}}$), the force-dependent lifetime $\tau(f)$ is co-determined by both the shear- and unzip-stretch regions. The significantly increased lifetime at the lower force range is a direct result of the increase in energy cost of breaking each base pair, the switched energy barrier, and the longer distance of diffusion.

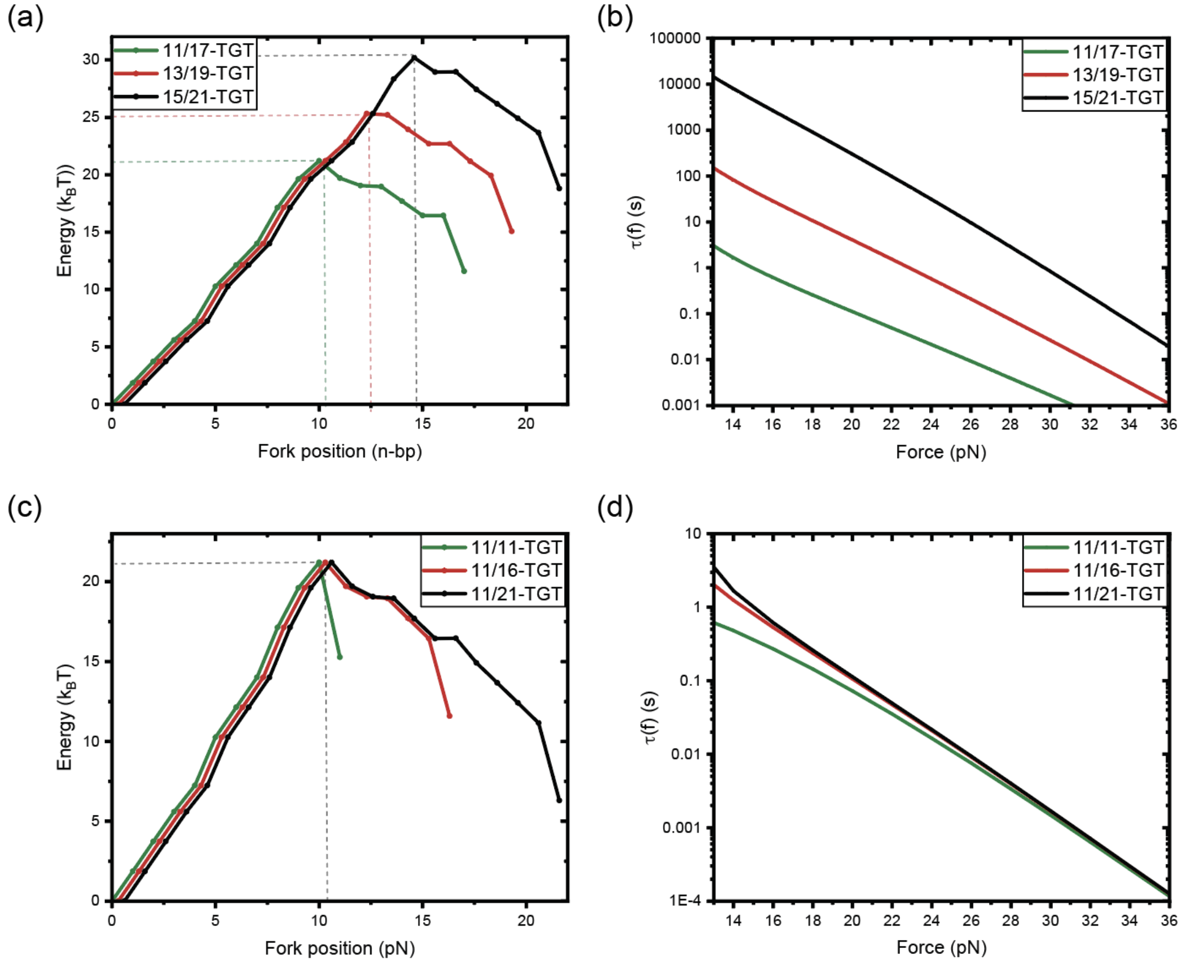


FIG. S1. Differential contributions to $\tau(f)$ from shear- and unzip-stretch regions at $f > f_c^u$. (a) The energy landscapes at 20 pN and (b) the tension-dependent lifetimes at high force above 18 pN of the 11/17-, 13/19-, and 15/21-TGTs with 10-, 12-, and 14-bp in shear-stretch region, respectively. (c) The energy landscapes at 20 pN and (d) the tension-dependent lifetimes at high force above 18 pN of the 11/11-, 11/16-, and 11/21-TGTs with 1-, 6-, and 11-bp in unzip-stretch region, respectively.

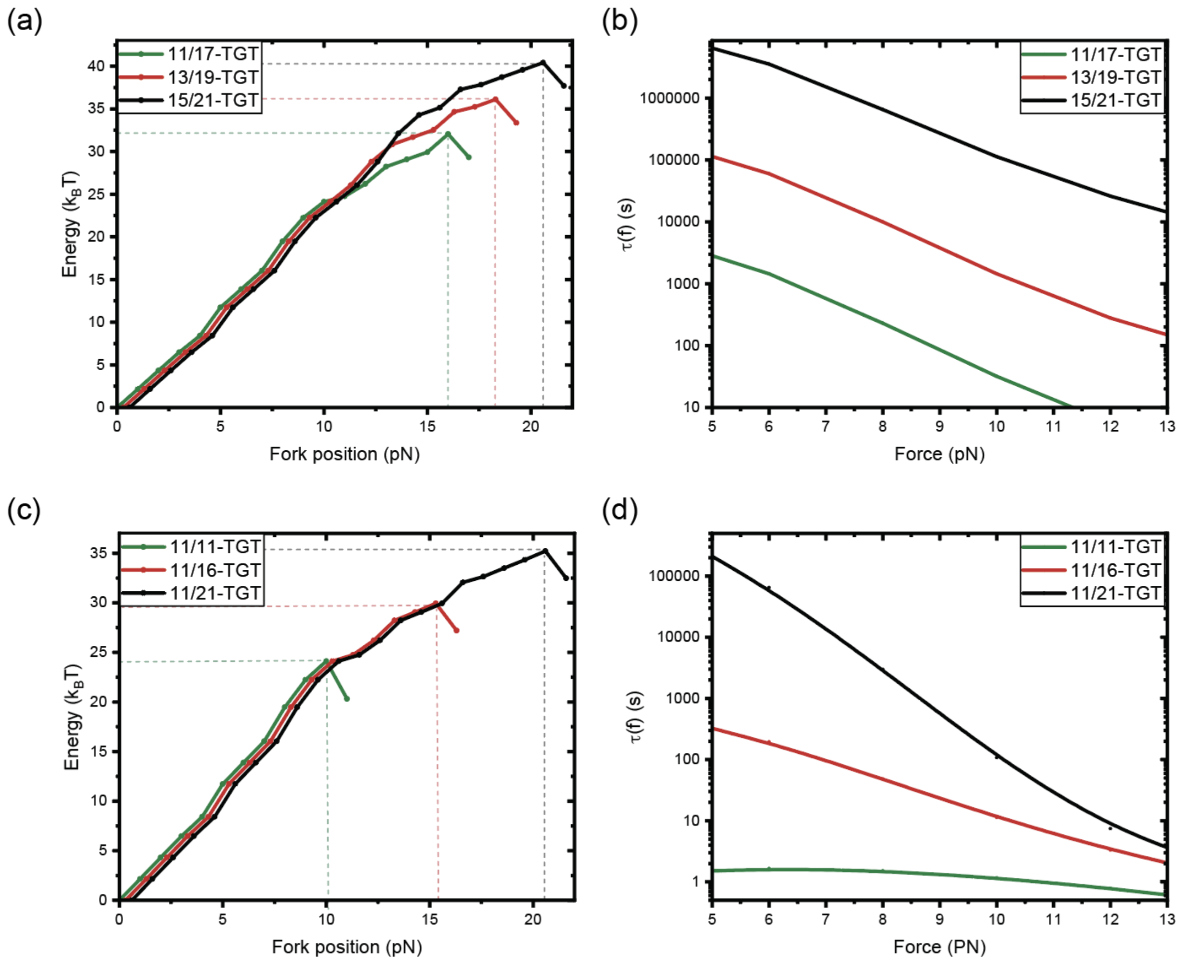


FIG. S2. Differential contributions to $\tau(f)$ from shear- and unzip-stretch regions at $f < f_c^u$. (a) The energy landscapes at 10 pN and (b) the tension-dependent lifetimes at high force below 18 pN of the 11/17-, 13/19-, and 15/21-TGTs with 10-, 12-, and 14-bp in shear-stretch region, respectively. (c) The energy landscapes at 10 pN and (d) the tension-dependent lifetimes at high force below 18 pN of the 11/11-, 11/16-, and 11/21-TGTs with 1-, 6-, and 11-bp in unzip-stretch region, respectively.
